# Fast haplotype matching in very large cohorts using the Li and Stephens model

**DOI:** 10.1101/048280

**Authors:** Gerton Lunter

## Abstract

The Li and Stephens model, which approximates the coalescent describing the pattern of variation in a population, underpins a range of key tools and results in genetics. Although highly efficient compared to the coalescent, standard implementations of this model still cannot deal with the very large reference cohorts that are starting to become available, and practical implementations use heuristics to achieve reasonable runtimes. Here I describe a new, exact algorithm (“fastLS”) that implements the Li and Stephens model and achieves runtimes independent of the size of the reference cohort. Key to achieving this runtime is the use of the Burrows-Wheeler transform, allowing the algorithm to efficiently identify partial haplotype matches across a cohort. I show that the proposed data structure is very similar to, and generalizes, Durbin’s positional Burrows-Wheeler transform.

## 1 Introduction

The genetic variation in a population of interbreeding individuals is highly structured, and the canonical model that describes this structure mathematically is the Kingman’s coalescent [1], later extended to also include recombination [2, 3]. Although mathematically elegant, it is challenging to use these models directly for statistical inference. In 2003, Li and Stephens introduced a model (LS) that both a good approximation of Griffith’s ARG, and computationally tractable [4]. As a result, LS now underpins a large range of key tools and scientific findings [5, 6, 7, 8]. Depending on whether the input sequence is haploid or diploid, LS in its straightforward implementation as a hidden Markov model (HMM) runs in linear or quadratic time in the number of reference haplotypes. While this is orders of magnitude more efficient than algorithms based directly on Kingman’s coalescent or the ARG, the recent availability of affordable DNA sequencing technology has provided access to very large reference sets, on which even the LS model is intractable in its standard implementation, so that current implementations of LS use heuristics to cope with data sets encountered in practice [7].

A very different algorithm that is making an impact in genomics is the BurrowsWheeler transform (BWT). Invented in 1994 [9], this transformation permutes an arbitrary text in such a way that the original text can be recovered, while at the same time improving the compressibility of the transformed text by increasing simple repetitions. In addition, the transformed text, even in compressed form, serves as an index that allows rapid searching in the original text. In genomics this idea has so far been used mainly for fast alignment of short reads against a large and relatively repetitive reference genome [10]. More recently, Durbin introduced a variant of the BWT, termed the Positional Burrows-Wheeler Transform (PBWT), that exploits the additional structure that exists in a set of haplotypes in a population sample [11]. These data, which are usually encoded as a series of 0s and 1s representing the absence or presence in a sample of particular genetic variants along a reference sequence, have a natural representation as a matrix, where rows represent samples and columns represent the particular positions in a reference. Local matches between samples are only relevant at matching positions, and exploiting this restriction leads to improvements over a standard application of the BWT. The resulting data structure again allows for fast haplotype searches against a database, and achieves very high compression ratios [11].

There are two main results in this paper. First, I establish a formal connection between the standard and positional BWT, showing how the PBWT as introduced in [11]is a special case of the BWT. This connection also shows how the PBWT can be slightly generalized to cope with the multiallelic case. Besides providing an additional perspective on the positional BWT algorithms, which helps to better understand them, it also provides a mechanical way to “lift” existing algorithms operating on the BWT data structure to their positional equivalent, allowing the large literature on BWT algorithms to be applied to the current data structure. I show how this works by deriving the haplotype search algorithm from the equivalent BWT algorithm.

The second contribution consist of algorithms that implement Li and Stephens’ model on top of the BWT. More precisely, I present algorithms that compute maximumlikelihood (“Viterbi”) paths through the Li and Stephens hidden Markov model, providing a parsimonious description of a given sequence as an imperfect mosaic of reference haplotypes. The ability to efficiently identify matches in the database of reference haplotypes result in considerable improvements in runtime over the standard implementation, reducing the linear and quadratic asymptotic runtime to empirical constant time, independent of the number of reference haplotypes. More precisely, for *H* samples of *n* loci each, the standard implementation runs in *O*(*Hn*) time for a haploid input sequence, and *O*(*H*^2^*n*) for a diploid input sequence, while the proposed algorithms run in empirical *O*(*n*) time in both cases. This allows the Li and Stephens model to be used on very large reference panels, without recourse to approximations.

## 2 Haplotype matching using the BWT

Let *x*_0_,…,*x*_*H*‒1_ be *H* haplotype sequences, each consisting of *n* symbols from the alphabet A representing the possible allelic states at a locus; for simplicity I will often use *A* = {0,1} in this paper. A straightforward way of identifying haplotype matches would be to use the BWT on the concatenation *x*_0_*x*_1_ … *x*_*H*⋯1_ of haplotype sequences. It turns out that a more efficient algorithm is obtained, in terms of time and memory use, by embedding this sequence of *Hn* characters into a sequence of 2*Hn* characters taken from a much larger alphabet. The increase in sequence length and alphabet size is offset by the additional structure in the BWT that results from the chosen embedding and additional characters. This in turn translates into better compression and a streamlined search algorithm.

### Algorithm 1 Calculating BWT(X)

**Input:** sequences *x*_0_,…, *x*_*H*‒1_, each of length *n*; alphabet *A*

**Output:** Block permutations 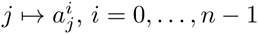.

1: 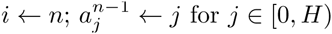
2: While *i* > 0:
3: 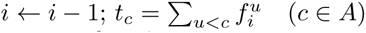
4: For *j* in [0, *H*):
5: 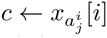
6: 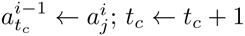

**Input:** sequences *x*_0_,…; *x*_*H*‒1_, each of length *n*; alphabet *A* = {0;1}

**Output:** Block permutations 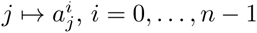

1: 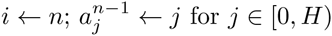
2: While *i* > 0:
3: 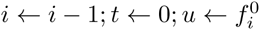
4: For *j* in [0;*H*):
5: 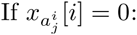
6: 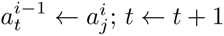
7: Else:
8: 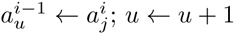

I will write *x*[*j*] for the *j*th symbol in the sequence *x*, and *x*[*j*, *k*) for the subsequence starting at position *j* and ending at *k* – 1. I will also use [*i,j*) to denote the halfopen interval {*i*, *i* + 1,…,*j* – 1}, and if *M*_*ij*_ is a matrix, *M*_*k*_ [*i*,*j*) is the subsequence *M*_*k*,*i*,_ *M*_*k*,*i*+1_,…, *M*_*k*,*j*‒1_ of the *k*th row of the matrix. Throughout this paper, all indices start at 0.

Let **p**_0_,…, **p**_*n*‒1_ be *n* additional symbols in the alphabet, ordered such that **p**_0_ < … < **P**_*n*‒1_ < 0 < 1. Introduce a new sequence *X* of length 2*Hn* by inserting a symbol **p**_*j*_ after each symbol *X*_*i*_[*j*] and concatenating the resulting sequences into a single sequence of the form

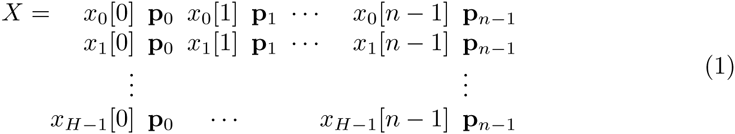

(To impose a particular initial ordering I will later on replace the last symbol **p**_*n*‒1_ by *H* symbols 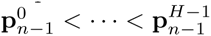, but to avoid cluttering the notation I ignore this detail for now.) Consider all cyclic shifts *X*^*k*^ = *X*[*k*]*X*[*k* + 1] … *X*[2*Hn* – 1]*X*[0] … *X*[*k* – 1] of *X*. Let *M* be the matrix obtained by writing *X*^*k*^ on the *k*th row of a square matrix, and sorting the resulting rows lexicographically. Let *π* be the permutation that sorts the rows, so that *X*^*π*(0)^ < *X*^*π*(1)^ < … < *X*^*π*(2*Hn*–1)^, and *M*_*ij*_ = *X*^*π*(*i*)^[*j*]. The BurrowsWheeler transform of *X* is the last column of this matrix: *BWT*(*X*)[*i*] = *X*^*π*(*i*)^[2*Hn* – 1]. Note that this is almost the traditional BWT of the sequence *X* – the difference is that there is no special ’end’ character. This character is used to identify the start of the sequence; here, the special structure of *X* is sufficient to navigate *BWT*(*X*).

Now consider how the matrix *M* may be constructed. The position symbols **p**_*i*_ determine the coarse structure of *M*, which is independent of the data *x*_*i*_ apart from the haplotype frequencies 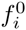 and 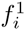 (see Figure 1). The fine-scale structure of *M* within each “block” of *H* rows is determined by the data. More precisely, rows in the block starting at index *iH* are those cyclic shifts of *X* that start with symbol **p**_*i*_ and end with *x*_*k*_[*i*] for some *k* ϵ [0, *H*), such that these rows are ordered lexicographically within the block. Let 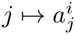 denote the permutation of [0, *H*) that describes this order within block i, so that row *iH* + *j* ends with symbol 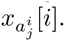. Determining *M* therefore boils down to determining the *n* permutations 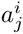 for *i* ∈ [0,*n*), since these determine the top half of *M*, and the remaining rows follow trivially from that (again, see Figure 1).

**Figure 1:**
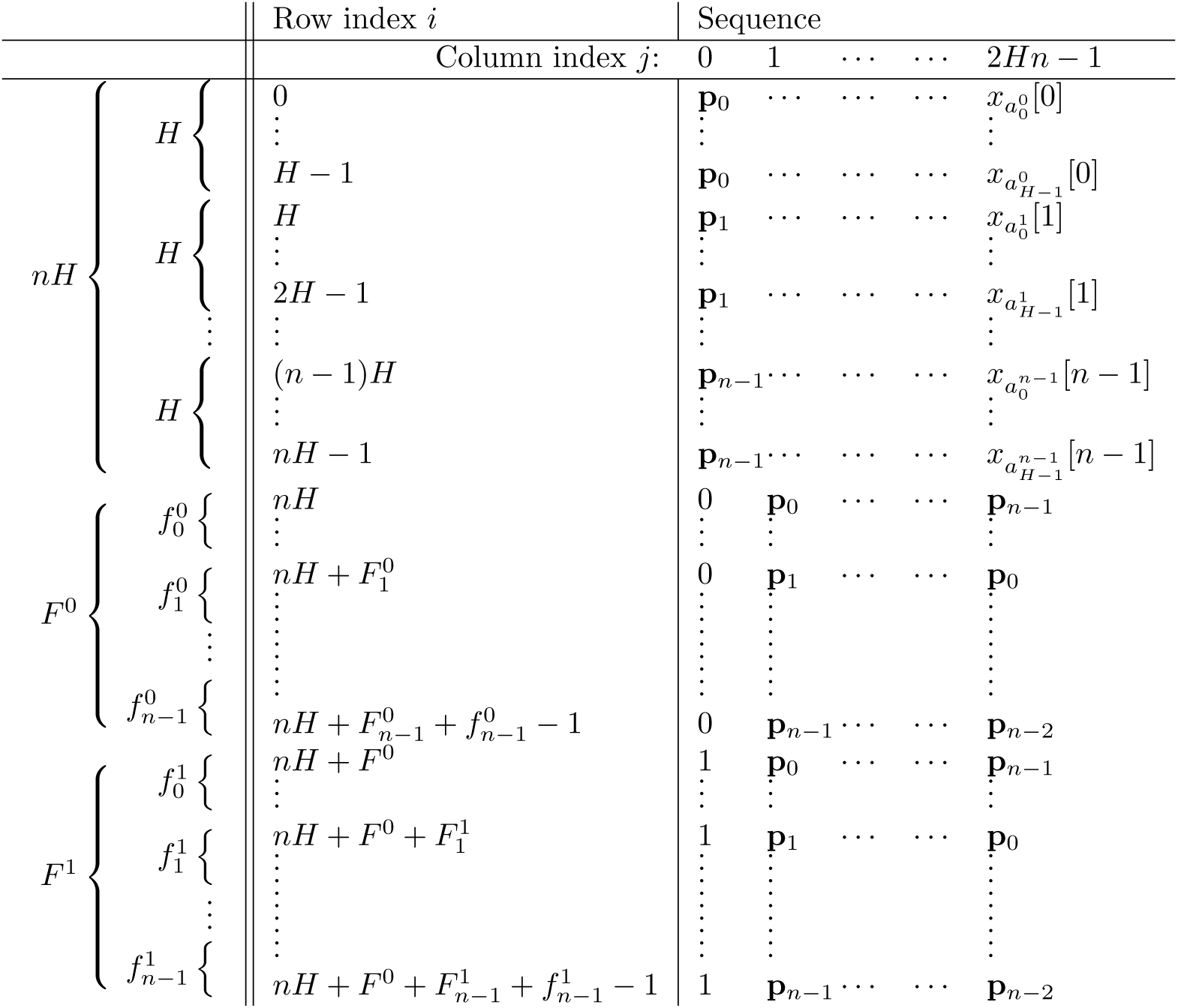
Structure of the matrix *M*_*ij*_. The rows *M*_*i*_ are sorted lexicographically; in particular **p** < **p**_1_ ⋯ < 0 < 1. The Burrows-Wheeler transform of *X* (see text) is the rightmost column of *M*, while the positional *BWT* of the sequences *x*_0_, ⋯,*x*_*H* – 1_ is the upper half of the same column (see text). The column indices are determined by 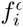, the allele frequency of symbol a at locus *i*, and 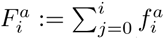, the cumulative frequency of symbol a across loci 0, ⋯, *i*. Note that ordering of rows (*n* – 1)*H* to *Hn* – 1 is determined by the special position symbols 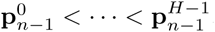, but to avoid cluttering the notation these are all written as **p**_*n*‒1_.

The permutations 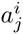 are determined recursively, working from *i* = *n* – 1 backwards. Because we imposed the special ordering 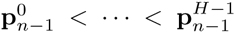 on the final position symbols, the permutation for block *n* – 1 is given by the identity permutation 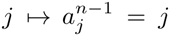. Now suppose the permutation 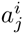 for block *i* has been determined. The sequences in block *i* – 1 are formed from those in block *i* by moving two characters from the end to the front. The first character in any sequence of this new block is *p*_*i*–1_, which does not influence the ordering within the block. The second character is an allele marker 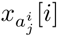. To sort the sequences in block *i* – 1 in lexicographic order, it is therefore sufficient to list those sequences that start a 0 symbol first, followed by those starting with a 1 symbol (followed by other symbols if the locus is multiallelic), and otherwise leave the original order undisturbed. Doing this results in Algorithm 1.

To show that the proposed construction is equivalent to the positional BurrowsWheeler transform, Algorithm 1 is given both for general alphabets A and specialized for the case *A* = {0,1}, since that in that case the inner loop is precisely Algorithm 1 in [11] (except that the proposed algorithm runs back-to-front, as is usual for BWT algorithms). As in the PBWT algorithm, the permutations 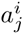 play the role of the suffix array in the ordinary *BWT* algorithm. Note that the output includes a permutation 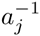, which encodes how the very first characters *x*_*j*_ [0] influence the permutation of the cyclic shifts *X*^*k*^; this permutation is used in Algorithm 5 below.

Following [11] I now define the PBWT of *x*_0_,…, *x*_*H*‒1_ as the first half of *BWT*(*X*), which is availably implicitly as 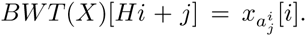. Figure 1 shows that the second half of *BWT* (*X*) is determined by the allele frequencies 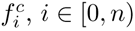, which can be computed easily from the relevant block in the first half of *BWT*(*X*), so that the PBWT of xo,…,*x*_*H*‒1_ is in fact equivalent to *BWT*(*X*).

## 3 Substring searching

Algorithm 1 to calculate *BWT*(*X*) in linear time makes use of the special structure of *x*, and is not a specialization of an existing, general algorithm to calculate the BWT. By contrast, Algorithm 2, which performs a substring search, can be derived directly from its analogous algorithm for a general BWT.

To describe the algorithm, let *M* be the sorted matrix of cyclic shifts of an arbitrary sequence *X* of length *n*, so that *BWT*(*X*)[*i*] = *M*_*i*_[*n* – 1], and let *R*^*a*^(*i*) (the “a-rank” for row *i*) be the number of times that a appears in *BWT*(*X*)[0, *i*). This function can be calculated from *BWT*(*X*) in *O*(*n*) time, and can be reduced to *O*(1) time using a small additional data structure that stores *R*^*a*^(*i*) for *i* values at regular intervals. Finally, let *C*(*a*) (the cumulative symbol frequency) be the number of symbols in *X* that are less than *a*. This notation makes it possible to write down Algorithm 2, for substring searching.

### Algorithm 2 General subsequence search

**Input:** sequences *w*[0,*j*) BWT(*X*) of sequence *X*[0; *n*)

**Output:** Indices *s*; *e* such that *M*_*k*_[0; *j*) = *w* for *k* ∈ [*s*; *e*)

1: *s* ← 0, *e* ← *n*, *i* ← *j*
2: While *s* < *e* and *i* > 0: ▹ *w*[*i*, *j*) matches *M*_*k*_[0, *j* – *i*)] for *k* ∈ [*s*, *e*)
3: *i* ← *i* – 1
4: *s* ← *C*(*w*[*i*]) + *R*^*w*[*i*]^ (*s*)
5: *e* ← *C*(*w*[*i*]) + *R*^*w*[*i*]^ (*e*)

To understand the algorithm, consider all rows of *M* that end with a symbol a. If these rows are cyclically shifted rightward, so that the last symbol becomes the first and all others are moved one position to the right, all rows will now start with a, and the relative order in which they appear in *M* (which they must as *M* contains all cyclic shifts of *X*) is the same as before the shift since they were ordered lexicographically to start with. Suppose that *M*_*k*_ is a row that ends with *a*, and that after right-shifting it ends up as row *M*_*k*′_; then the above observation means that the rank *R*^*a*^(*k*) of the symbol *a* in *M*_*k*_ in the last column of *M*, is the same as the rank in the first column of *M* of the symbol *a* in *M*_*K*′_. Because *M* is sorted lexicographically, the rows that start with *a* form *a* contiguous block in *M*, so that the first-column rank of the symbol *a* in row *M*_*k*′_ is *k* – *C*(*a*), so that *R*^*a*^(*k*) = *k′* – *C*(*a*) or

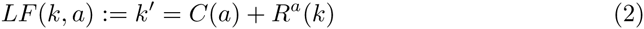

The function *k* ↦ *LF*(*k*, *a*), mapping row *k* to the row corresponding to its right-shifted counterpart *k*′, is called the *last-to-first* mapping because it maps the last (rightmost) symbol of *M*_*k*_ to the corresponding symbol in the first (leftmost) position of *M*_*k*′_. It is repeatedly used to identify the interval of rows corresponding to sequences that match one additional character of *w*.

Note that the mapping is well-defined whether or not *M*_*k*_[*n* – 1] = *a*. This makes it possible to think of *k* as representing a possible location between two entries (*k* and *k* – 1) in *M* where a sequence (or sequence prefix) *x* not necessarily represented in *M* would be inserted; this is the view taken in the search algorithm. Alternatively, when *k* is thought of as a particular row in *M*, that row’s initial character *a* can be obtained from the *C*(·) function, and since the mapping (2) is invertible when restricted to the set of rows *k* ending in *a*, this makes the mapping *k* ↦ *LF*(*k*, *M*_*k*_ [*n* – 1]) invertible for all *k*. The existence of this inverse mapping also follows directly from the observation that it corresponds to rotating the sequence one position leftward; it could be called the *first-to-last mapping, k* ↦ *FL*(*k*), and is used in Algorithm 5 below.

To derive the corresponding algorithm for matching a sequence in the PBWT data structure, it is enough to track the bounding variables for two steps through the standard BWT algorithm acting on the “lifted” sequence *X*, matching a haplotype character and a position character. The first step identifies the new range depending on the haplotype character to be matched, and points these variables to the second half of the matrix. The next step moves the bounding variable back into the first half by moving a position character in front. Because of the regular form of *BWT*(*X*) (see Figure 1), these two steps can be followed algebraically and combined into a single update step. The derivation, which is straightforward but requires additional notation, is presented in the Appendix. The resulting combined update step is given by a modified last-to-first mapping function, which now additionally depends on the current position *i*:

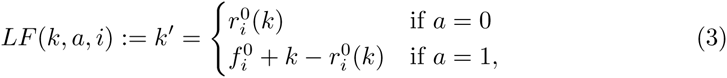

or for an arbitrary alphabet, 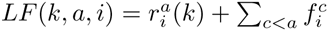.Here 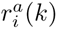 is the posiional analogue of *R*^*a*^(*i*), and counts how often *a* appears in the first *k* rows of the *i*th block of *PBWT*(*x*_0_,…, *x*_*H–1*_), or equivalently, in 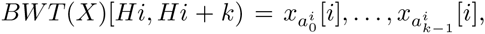,and 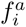 is the (haplotype) frequency of *a* at position *i*. This leads to Algorithm 3.

### Algorithm 3 PBWT subsequence search

**Input:** Sequence *w*[0, *j*), PBWT of *x*_0_,…, *x*_*H* – 1_

**Output:** Indices *s*, *e* such that 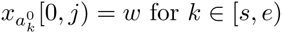

1: *s* ← 0, *e* ← *H*, *i* ← *j*
2: While *s* < *e* and *i* > 0: ▹ *w*[*i*, *j*) matche 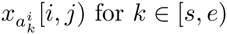
3: *i* ← *i* – 1
4: *s* ← *LF*(*s*, *w*[*i*], *i*) ▹ see equation (3)
5: *e* ← *LF*(*e*, *w*[*i*], *i*)

## 4 Haploid Li and Stephens

The Li and Stephens (LS) model approximates the coalescent model describing the relationship between DNA sequences in a population, by generating a new sequence as a mosaic of imperfect copies of existing sequences [4]. The popularity of the model stems from the fact that it is both a good approximation to the full coalescent model with recombination, as well as fast to compute in its natural implementation as a hidden Markov model, running in *O*(*Hn*) time for *H* sequences of length *n*. however, for very large population samples this is still too slow in practice.

Here I describe an algorithm to compute the maximum likelihood path through the LS hidden Markov model (HMM) in empirical *O*(*n*) time. The approach is not to consider single sequences to copy fro*M*, but *groups* of sequences that share a common subsequence. Like the Viterbi algorithm for hMMs, the proposed algorithm traverses the sequence to be explained, but rather than using a dynamic programming approach, it uses a branch-and-bound approach considering (groups of) potential path prefixes to a maximum likelihood path. Where at each iteration the Viterbi algorithm must consider all possible sequences that a potential path prefix could end with, the proposed algorithm in principle considers all extensions of the current potential path prefixes (the “branch”part), but ignores prefixes that cannot be part of an optimal path (the “bound” part). For instance, if a prefix can be extended with a matching nucleotide, a recombination does not have to be considered, since the recombination can be postponed at no cost. Below I will show this more formally. This formal approach is perhaps not necessary (or even helpful) for the haploid case, but becomes useful when *i* introduce the diploid Li and Stephens algorithm.

### Algorithm 4 Haploid Burrows-Wheeler Li and Stephens

**Input:** Sequence *w*[0, *n*), PBWT of *x*_0_,…, *x*_*H*‒1_ scores *μ* ≥ 0, *ρ* ≥ 0.

**Output:** Minimum path score under the Li and Stephens model

1: *i* ← *n*; *st* ← [(0, *H*, 0)]; *gm* ← 0; *traceback* ← [(*n* – 1, –1, –1)]
2: While *i* > 0: ▹ *st* represent states of paths in a full suffix set for *x*[*i*, *n*)
3: *i* ← *i* – 1; *st*′ ← []; *gm*′ ← *gm* + *μ*; *extended* ← *False*
4: For (*s*,*e*, *score*) in *st*:
5: If *score* < *gm* + *ρ*:
6: *s*′ ← *LF*(*s*, *x*[*i*], *i*); *e*′ ← *LF*(*e*, *x*[*i*], *i*)
7: If *s*′ < *e*′:
8: *st*′.*append*((*s*′, *e*′, *score*))
9: *gm*′ ← min(*gm*′, *score*)
10: If *score* = *gm*: extended ← True
11: If *score* + *μ* < *gm*′ + *ρ*:
12: *s*′ ← *LF*(*s*, 1 – *x*[*i*], *i*); *e*′ ← *LF*(e, 1 – *x*[*i*], *i*)
13: If *s*′ < *e*′: *st*′.*append*((*s*′, *e*′, *score* + *μ*))
14: *s*′ ← *LF*(0, *x*[*i*], *i*); *e*′ ← *LF*(*H*, *x*[*i*], *i*)
15: If *s*′ < *e*′ and *extended* = *False*: ▹ Never true on 1st iteration
16: *st*′.*append*((*s*′, *e*′, *gm* + *ρ*))
17: *traceback.append*((*i*, *gm*_*idx*, *gm* + *ρ*))
18: *gm* ← *gm*′; *st* ← *st*′
19: *gm*_*idx* ← any of {*s*|(*s*, *e*, *score*) ∈ *st* and *score* = *gm*}
20: Return *gm*, *gm*_*idx*, *traceback*

First some definitions. A *placed character* is a character c at a sequence position *i*; it is equivalent to a pair *c***p**_*i*_ where **p**_*i*_ is the position symbol introduced before. Two placed characters are *contiguous* if they occupy neighbouring positions; subsequences of placed characters are contiguous if every pair of neighbouring characters is; and two or more subsequences are contiguous if their concatenation is. A *path π* of *M parts* through a set of sequences Ω = {*x*_0_,…, *x*_*H*‒1_} is a contiguous sequence of *M* subsequences *s*_0_,…, *s*_*m*‒1_ such that each *s*_*i*_ is a subsequence of some *x*_*j*_. I will write a path as

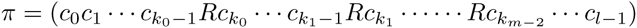

where *c*_*i*_ is a character placed at position *i*, and *k*_0_, *k*_1_,…, *k*_*m*–2_ are the *recombination breakpoints* identified by the symbol *R* (which is not part of the alphabet), and *l* is the *length* of the path. The (sequence) *group* associated with *π* is the set *G*(*π*) of all sequences *X* ∈ Ω for which the subsequences *x*[*k*_*m*–2_, *l*) agree with the suffix *c*_*k*_*m*–2_ · · · *c*_*l*_–1_ that follows the last recombination in *π*. The *extension πc*_*l*_ (of length *l* + 1) is the path ( ⋯ *Rc*_*km*-2_ ⋯ *c*_*l*-1_*c*_*l*_;), if it exists; since by definition all subsequences that make up a path are subsequences of some *x*_*j*_, existence of an extension implies that its group is nonempty. The extension *πR* (of length *l*) is defined as ( ⋯ *Rc*_*km*–2_ ⋯ *c*_*l*‒1_*R*), and always exists; its group is Ω. Finally, the *path prefix π*[0,*t*) is the path (*c*_0_ ⋯ *c*_*t*‒1_) including any *R* symbols for recombinations between positions 0 and *t* – 1; a path prefix never ends with an *R* symbol.

### Algorithm 5 Haploid traceback

**Input:** Sequence *x*[0, *n*), PBWT of *x*_0_,…, *x*_*H*-1_, scores *μ* ≥ 0, *ρ* ≥ 0, minimum *score gm*, corresponding index *gm*_*idx*, traceback list *traceback*.

**Output:** Representation path of a minimum-scoring path

1: Function *FL*(*k*, *i*): ▹ “First-to-last” mapping
2: 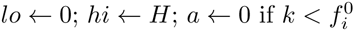 else 1
3: While *lo* < *hi*: ▹ *LF*(*j*, *a*, *i*) ≤ *k* ∀_*j*_ < *lo* and *LF*(*j*, *a*, *i*) > *k* ∀_*j*_ > *hi*
4: *mid* ← ⎿(*lo* + *hi*)/2⏌
5: If *LF*(*mid*, *a*, *i*) ≤ *k*: *lo* ← *mid* + 1
6: Else: *hi* ← *mid*
7: Return *a*, *lo* – 1
8: 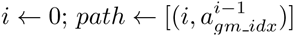
9: For (*t*_*locus*, *t*_*idx*, *t*_*score*) in reverse(*traceback*):
10: While *i* ≤ *t*_*jocus*:
11: *a*, *gm*_*idx* ← *FL*(*gm*_*idx*, *i*)
12: If a ≠ *x*[*i*]: *gm* ← *gm* ← ←
13: *i* ← *i* + 1
14: If *gm* = *t*_*score*:
15: *gm*_*idx* ← *t*_*idx*; *gm* ← *gm* – *ρ*; 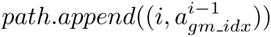
16: Return *path*

For a given sequence *X* and a path *π*, the Li and Stephens model assigns a joint likelihood to the event that *π* occurred and gave rise to sequence *x*. If *π* has *M* parts and has *k* mismatches to *x*, this likelihood is

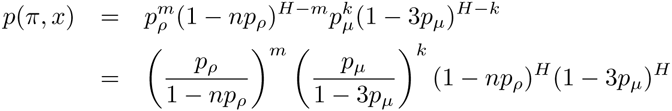

where *p*_*ρ*_ is the probability of recombining into a particular other sequence, and *p*_*μ*_ is the probability of a mutation to one of the three other nucleotides. The negative log likelihood takes a particularly simple form,

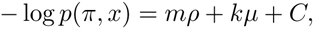

where *C* is a constant, *ρ* = – log(*p*_*ρ*_/(1 – *np*_*ρ*_)) and *μ* = – log(*p*_*μ*_/(1 – 3*p*_*μ*_)). This motivates defining the *path score* as *s*_*x*_(*π*) = *mρ* + *kμ*,where *m* and *k* are defined as above. *i* drop the subscript *x* from *s*_*x*_(*π*) when this is possible without creating confusion.

Suppose we want to calculate a path *π* that minimizes *s*(*π*). This can be done by iteratively constructing path prefixes *π*′,so that at each step one of them is a prefix of a full path *π* that minimizes *s*(*π*). Note that the minimum score achievable by a path *π* that has *π*′ as its prefix depends on the prefix score *s*(*π*′) and the prefix group *G*(*π*′), but not on the rest of the prefix. This is because *G*(*π*′) is the set of sequences the Li and Stephens model could be copying from at the end of *π*′, and the Markov property of the model implies that the minimum score only depends on the sequence being copied from (and the prefix score). This justifies the definition of *state* of a path (prefix) *π*′ to be the pair (*G*(*π*′), *s*(*π*′)).

The key observation for the algorithm is that some states (*G*, *s*) can be ignored, because any of their extensions give rise to paths and scores that are also achievable via other states. To make this precise I need one more definition. A set *S* of path prefixes, all of length *l*, is a *full prefix set* for *x*[0, *l*) if for any sequence *x*′ whose prefix *x*′[0, *l*) agrees with *x*[0, *l*), there exists a path *π* that achieves the minimum score (i.e. *s*_*x*′_ (*π*) = min_*π*′_ *s*_*x*′_ (*π*′)) and whose prefix *π*[0, *l*) is in *S*. If we can somehow find a way to iteratively construct full prefix sets of increasing length, the problem of finding a minimum-score path is solved, because the required path will be an element of the full prefix set for the full-length sequence *x*. The following theorem shows how to do this:

### Theorem 1.

Suppose S is a full prefix set for x[0, l), S′ a set of prefixes of length l + 1, and let s_mjn_ = min_π ∈S_s (π) and 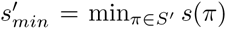 = min_π∈S′_ s(π). Then S′ is a full prefix set for x[0, l + 1) if the following two conditions are true:

a For all π ∈ S and all a ∈ {0, 1} so that πa is an extension and s(πa) < 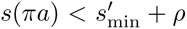 we have πa ∈ S′.
b If there is no π ∈ S so that s(π) = s_min_ and πx[l] is an extension, then S′ contains a path of the form πRx[l] with s(π) = s_min_.

In other words, certain extensions are *not* required to be in *S*′: extensions *πa* whose score exceed the minimum plus *ρ* can be left out (since a recombination from the minimum-scoring prefix would give a path that is at least as good), and recombinations can be ignored altogether as long as any current lowest-scoring path has a matching extension (since otherwise postponing the recombination would again be at least as good) – and if not, only a single recombination from a lowest-scoring path needs to be considered.

Algorithm 4 implements these ideas. It does not actually construct prefix sets of paths, but sets of *states* of paths in prefix sets. This is sufficient since the state determines how paths can be extended. By using the PBWT, these states can be represented efficiently, using just the score and a pair of indices into the PBWT that correspond to a set of subsequence matches to sequences in Ω, similar to how the variables *S* and e in Algorithm 3 represent the interval [*s*, *e*) corresponding to a set of subsequence matches. Another difference with the description above is that the algorithm scans the sequence back-to-front, extending partial matches leftward, so that the invariant refers to the full *suffix* set, rather than the full prefix set.

The algorithm computes *gm* = *s*_min_, and keeps a running minimum score *gm*′ that bounds 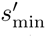, ignoring states whose new score are not less than *gm*′ + *ρ*. At the end of an iteration, states whose score are not lower than the now updated *gm*′ plus *ρ* are not immediately removed, but are instead ignored in the next iteration. The algorithm implicitly considers both score bounds implied by *gm* and *gm*^′^, but in each situation uses only the tighter bound of the two to decide which states to ignore.

It is possible for different paths to result in overlapping or identical states, resulting in duplicate or otherwise redundant entries in the *st* array. Although redundant entries do not impact the correctness of the algorithm, they can dramatically reduce efficiency. A practical implementation therefore includes a step that occasionally removes redundant states.

The algorithm can be generalized a little by allowing the mutation score *μ* ≥ 0 to depend on the position. The path score is then defined as *s*(*π*) = *mρ* + ∑_*i:x*[*i*]≠*π*[*i*]_*μ*_*i*_. Theorem 1 continues to hold, and so does Algorithm 4, with the obvious changes. The current approach does not lend itself easily to generalize to a position-dependent recombination probability, as the proof of Theorem 1 relies on delaying the recombination without changing the score, which is only possible if *ρ* is constant along the sequence.

Note that the algorithm can be simplified when *μ*_i_ ≥ 2*ρ*, because a mismatch can always be circumvented by two recombinations (before and after the offending locus), so that only exact matches need to be considered. In human genetics polymorphisms are sparse, and recombinations can only be localized to within hundreds or thousands of positions. Even when a maximum likelihood path is sought it is natural to marginalize over these positions, and this makes the probability of a recombination between two polymorphic sites at least an order of magnitude higher than the probability of a mutation, so that *μ* ≫ *ρ*. however, in the presence of phasing errors the probability of a mismatch can be much higher than that of a mutation, so that the regime *μ* < 2*ρ* is of practical importance.

Algorithm 4 only computes the optimal score, and to obtain an optimal-scoring path *π* itself a backtracking step is needed (Algorithm 5). here it is useful that Algorithm

4works in the backward direction, so that the result of the backtracking is oriented in the natural direction. To track an optimal path along a sequence, the PBWT index corresponding to that sequence can be tracked using the “first-to-last” mapping, inverting the steps in lines 6 and 12 in Algorithm 4, and the minimum score of the remaining suffix is updated whenever a difference between this sequence and *x* is found. Recombinations are followed greedily, as it is always correct to follow a feasible recombination, and it is never clear whether a particular recombination is the last feasible one for a particular sequence. Algorithm 4 collects information about recombinations in the *traceback* list, and when a recombination and score is identified that forms a feasible suffix to the path so far, it is followed.

The naive implementation of Algorithm 5 is somewhat slower than the haploid Li and Stephens algorithm itself, due to the *FL* function which takes *O*(log *H*) time in the implementation shown. In practice the *PBWT* will be stored in compressed form using run-length encoding, which allows a faster implementation of *FL*.

## 5 Diploid Li and Stephens

Where the haploid Li and Stephens algorithm computes a single haplotype path maximizing the probability of a given haploid sequence, the diploid Li and Stephens algorithm aims to find a *pair* of haplotype paths that maximizes the probability of a sequence of diploid *genotypes* under the same model. The approach used to derive the haploid algorithm also works in this case, but the details are more involved.

Let *x* be a sequence of genotypes, encoded as values 0, 1 or 2 at each position representing homozygous ancestral, heterozygous, and homozygous derived genotypes. The aim is to compute a pair of paths *α*, *β* that minimizes a score. As before this score contains terms for recombinations and mismatches, but the mismatch term now considers genotypes rather than haplotypes. More precisely, the score associated to the pair {*α*, *β*} is defined as *s*(*α*, *β*) = *ρm*(*α*) + *ρm*(*β*) + *μK*(*α*, *β*), where *m*(*α*) represents the number of parts of path *α*, as before, and *K* = ∑_*i*_ |*α*[*i*] + *β*[*i*] – *x*[*i*]| counts the number of mismatches of the paths *α* and *β* to the genotype sequence *x*.

The approach of the algorithm is similar to the haploid case, again sequentially building full prefix sets for ever longer sequence prefixes until a minimum path pair is found. To describe the approach, the definitions of sequence group, state and full prefix set need to be modified.

The *sequence group* associated to an unordered pair of paths {*α*, *β*} is defined as *G*(*α*, *β*) = {{*x*, *y*}|*x* ∈ *G*(*α*), *y* ∈ *G*(*β*)}. Similarly, using the same justification as before, the *state* of an (unordered) path pair {*α*, *β*} is defined to be the pair (*G*(*α*, *β*), *s*(*α*, *β*)). A *full prefix set S* for *x*[0, *l*) is defined as a set of (unordered) pairs of path prefixes such that for any sequence *x*′ that extends *x*[0, *l*), there exists a path pair {*α*, *β*} that achieves the minimum score *s*_*x*′_ (*α*, *β*) = min_*α*′_,_*β*′_ *s*_*x*′_ (*α*′, *β*′) and whose prefix pair {*α*[0, *l*), *β*[0, *l*)} is in *S*. Finally, to formulate the theorem it is handy to introduce the notation *S̅* to denote the set of “haplotype” paths in *S*, or formally *S̅* = {*α*|{*α*,*β*} ∈ *S*}.

### Theorem 2.

Suppose S is a full prefix set for x[0, l) and S′ is a set of prefixes of length l + 1. Let S_min_(α) = min_β:{α,β}∈S_ ^S^_x_(α, β), 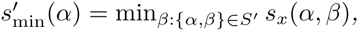 and s_min_ = min_α_S_min_ Sx(α, β), and s_m_in = min_a_ s_min_(α), 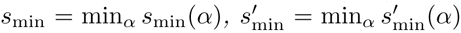. Then S′ is a full prefix set for x[0, l + 1) if the following are true:

a For all {α, β} ∈ S and a, b ∈ {0, 1}, so that αa and βb are both extensions and 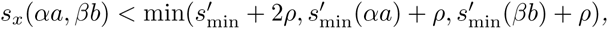, we have {αa, βb} ∈ S′.
b (If x[l] = 1:) For all α ∈ S̅ and a, b ∈ {0,1} with a + b =1, so that there is no β′ satisfying {α, β′} ∈ S and s(α, β′) = s_min_(α) and both αa and β′b are extensions, S′ contains a path pair of the form {αa, βRb} with {α, β} ∈ S and s(α, β) = s_min_(α).
c (If x[l] = 2b:) For all α ∈ S̅ and a ∈ {0,1}, so that there is no β′ satisfying {α, β′} ∈ S and s(α, β′) = s_min_(α) and both αa and β′b are extensions, S′ contains a path pair of the form {αa, βRb} with {α, β} ∈ S and s(α, β) = s_min_(α).
d (If x[1] = 2b:) If there is no pair {α′, β′} for which s(α′, β′) = s_min_ and either α′b or β′b is an extension, then S′ contains a path pair of the form {αRb, βRb} with {α, β} ∈ S.

Algorithm 6 implements these ideas. The core of the algorithm is formed by lines 11and 14 that consider regular extensions with a pair of characters (*a*_1_,*a*_2_); lines 2223 and 27 that consider single recombinations; and line 34 that considers simultaneous recombinations in both haplotypes. The remainder of the algorithm is concerned with implementing the conditions of Theorem 2 to ensure that redundant extensions are ignored. The variables *gm* and *gm*′ keep track of the current and next global minimum score *s*_min_ and 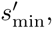, while the associative arrays *lm*[] and *1m*′[] keep track of *s*_min_(*α*) and 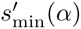 respectively. The associative array *extended*[] keeps track which paths *α* have a partner *β*′ that achieves the minimum score *s*_min_(*α*), and for which both *α* and *β*′ have extensions required in conditions *b* and *c*; whether the extension is appropriate is computed by the function *consider_recomb*. Finally, the variable *double_recomb* is used to ensure that at most one double recombination is considered at every iteration.

### Algorithm 6 Diploid Burrows-Wheeler Li and Stephens

**Input:** *x*[0, *n*) ∈ {0,1, 2}^*n*^, PBWT of *x*_0_,…, *x*_*H*-1_, *scores μ* ≥ 0, *ρ* ≥ 0.

**Output:** Minimum pair path *score* under the diploid Li and Stephens model

1: Function consider _recomb(*c*, *a*_1_, *a*_2_, *j*):
2: If *c* =1: Return (*a*_1_ + *a*_2_ = 1)
3: Else: Return (*a*_*j*_ = *c*/2)
4: *i* ← *n*; *st* ← [(0, *H*, 0, *H*, 0)]; *gm* ← 0; *lm*[(0, *H*)] ← 0;
5: *traceback* ← [(*n* –1, –1, –1, –1, –1)]
6: While *i* > 0: ▹ *st* represent states of path pairs in a full suffix set for *x*[*i*, *n*)
7: *i* ← *i* – 1; *st*′ ← []; *gm*′ ← *gm* + 2*μ*; *lm*′ ← {}; *extended* ← {}
8: *double_recomb* ← *False*
9: For (*s*_1_, *e*_1_, *s*_2_, *e*_2_, *score*) < (*a*_1_, *a*_2_) in *st* × {0,1} < {0,1}:
10: *score*′ ← *score* + *μ*|*a*_1_ + *a*_2_ – *x*[*i*]|
11: 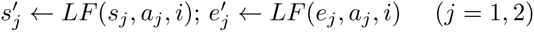
12: If 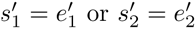 or *s*_2_ = *e*_2_ or (*s*_1_ = *s*_2_ and *e*_1_ = *e*_2_ and *a*_1_ > *a*_2_) or *score* ≥ min(*lm*[(*s*_1_, *e*_1_)] + *ρ*, *lm*[(*s*_2_, *e*_2_)] + *ρ*, *gm* + 2*ρ*) or 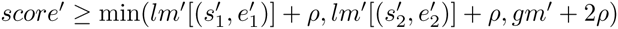:
13: continue
14: 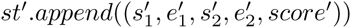
15: 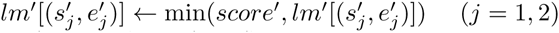
16: *gm*′ ← min(*score*′, *gm*′)
17: If *consider_recomb*(*x*[*i*], *a*_1_, *a*_2_, *j*) and *score* = *lm*[(*s*_3–*j*_, *e*_3–*j*_)]:
18: extended.insert((*s*_3–*j*_, *e*_3–*j*_, *a*_3–*j*_)) (*j* = 1, 2)
19: For (*s*_1_, *e*_1_, *s*_2_, *e*_2_, *score*) < (*a*_1_, *a*_2_, *j*) in *st* × {0,1} × {0,1} × {1, 2}:
20: *a*_*r*_ ← *a*_*j*_; *a*_*x*_ ← *a*_3–*j*_; s_*x*_ ← s_3–*j*_; e_*x*_ ← s_3–*j*_
21: *score*′ ← *score* + *ρ* + *μ*|*a*_*r*_ + **a**_*x*_ – *x*[*i*]|
22: 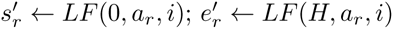
23: 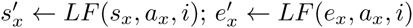
24: If *n*ot consider_recomb(*x*[*i*], *a*_1_, *a*_2_,*j*) or 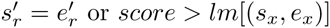 or *score* > *lm*[(*s*_*x*_, *e*_*x*_)] or (*s*_*x*_, *e*_*x*_, *a*_*x*_) ∈ extended or 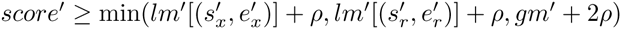:
25: continue
26: If 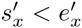:
27: 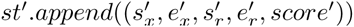
28: 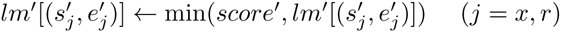
29: *gm*′ ← min(*score*′, *gm*′)
30: *extended.insert*((*s*_*x*_, *e*_*x*_, *a*_*x*_))
31: *traceback.append*((*i*, *s*_*x*_, *e*_*x*_, *s*_r_, *score* + *ρ*)) ▹ Not *score*′!
32: If *x*[*i*] ≠ 1 and *x*[*i*] = *a*_r_ + *a*_*x*_ and not *doub1e*_*recomb* and *score* = *gm* and (*s*_*r*_, *e*_*r*_, *a*_*r*_) ∉ *extended*:
33: If 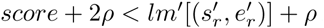
34: 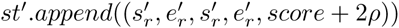
35: traceback.append((*i*, *s*_*x*_, –1, *s*_*r*_, *score* + 2*ρ*))
36: *doub1e_recomb* ← *True*
37: *gm* ← *gm*′; *lm*′ ← *lm*′; *st* ← *st*′
38: *gm*_*idx*1, *gm*_*idx*2 ← any of {*s*_1_, *s*_2_ |(*s*_1_, *e*_1_, *s*_2_, *e*_2_, *score*) ∈ *st* and *score* = *gm*}
39: Return *gm*, *gm*_*idx*1, *gm*_*idx*2, *traceback*

The traceback algorithm for diploid Li and Stephens is similar to the haploid algorithm. Again, the *traceback* list contains records describing the recombinations that have been considered. These records now additionally contain a pair *s*_*x*_, *e*_*x*_ that represent the range of PBWT indices corresponding to the sequence that does not undergo a recombination. As with the haploid algorith*m*, the traceback algorithm follows a recombination only if the path scores agree, but now also ensures that the index of the non-recombining path is contained in the range [*s*_*x*_,*e*_*x*_). Double recombinations are encoded by setting *e*_*x*_ = –1, and for such recombinations only the scores need to agree. A pseudocode implementation is given as Algorithm 7.

### Algorithm 7 Diploid traceback

**Input:** Sequence *x*[0, *n*), PBWT of *x*_0_,…, *x*_*H*-_1, *scores* ← ≥ 0, ≥ 0, minimum *score gm*, corresponding indices *gm*_*dxl*, *gm*_*idx*2, traceback list traceback.

**Output:** Representation path1, path2 of a minimum-scoring diploid path

1: Function *FL*(*k*, *i*): ▹ “First-to-last” mapping
2: *lo* ← 0; *hi* ← *H*; *a* ← 0 if 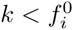 else 1
3: While *lo* < *hi*: ▹ *LF*(*j*, *a*, *i*) ≤ *k* ∀*j* < *lo* and *LF*(*j*, *a*, *i*) > *k* ∀*j* ≥ *hi*
4: *mid* ← ⎿(*lo* + *hi*)/2⏌
5: If *LF*(*mid*, *a*, *i*) ≤ *k* : *lo* ← *mid* + 1
6: Else: *hi* ← *mid*
7: Return *a*, *lo* – 1
8: 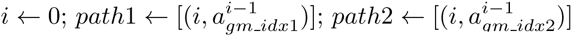
9: For (*t*_*locus*, *t*_*start*, *t*_*end*, *t*_*idx*, *t*_*score*) in reverse(traceback):
10: While *i* < *t*_*locus*:
11: *a*1, *gm*_*idx*1 ← *FL*(*gm*_*idx*1, *i*); *a*2,*gm*_*idx*2 ← *FL*(*gm*_*idx*2, *i*)
12: *gm* ← *gm* – *μ*|*a*1 + *a*2 – *x*[*i*]|; *i* ← *i* + 1
13: If *gm* = *t*_*score*:
14: If *t*_*end* = – 1: ▹ Double recombination
15: *gm*_*idx*1 ← *t*_*start*; *gm*_*idx*2 ← *t*_*idx*
16: 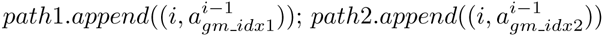
17: *gm* ← *gm* – 2*ρ*
18: Else If *t*_*start* ≤ *gm*_*idx*1 < *t*_*end*: ▹ Single recombination in path 2
19: 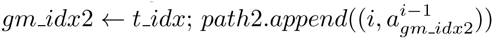
20: *gm* ← *gm* – *ρ*
21: Else If *t*_*start* ≤ *gm*_*idx*2 < *t*_*end*: ▹ Single recombination in path 1
22: 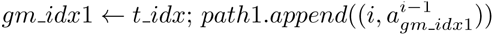
23: *gm* ← *gm* – *ρ*
24: Return *path*1, *path*2

**Figure 2:**
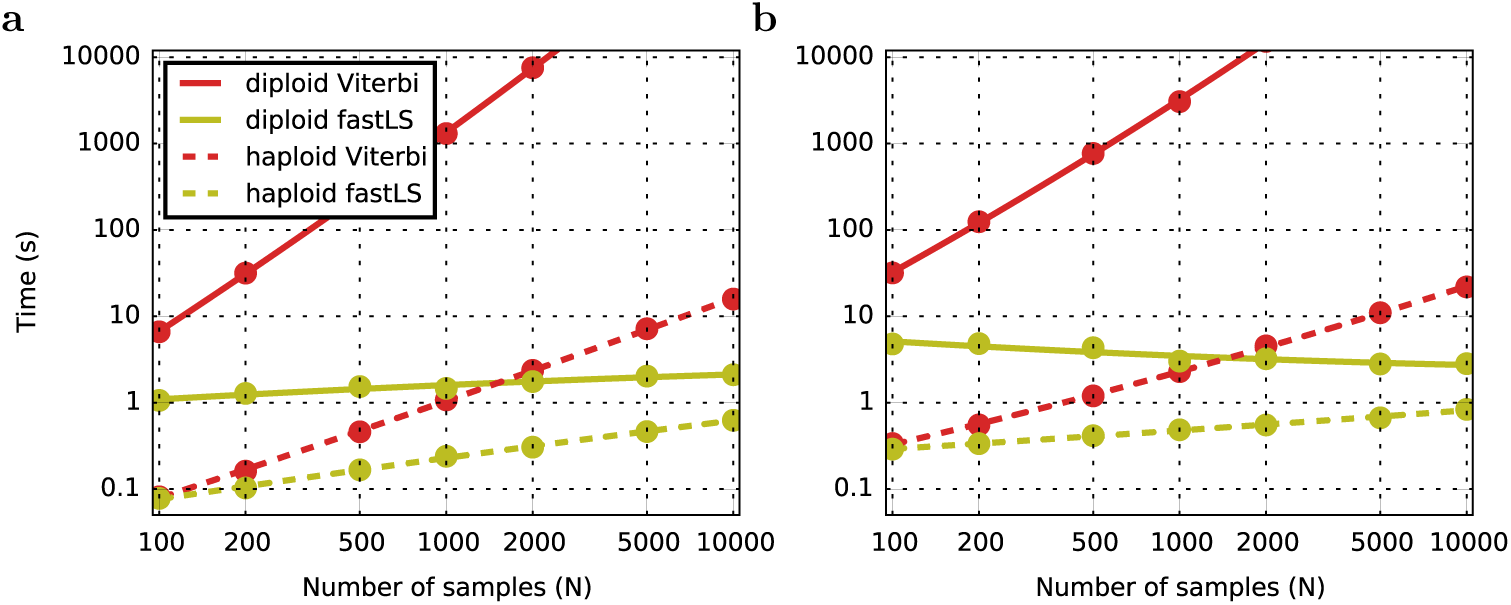
Running time for inferring inheritance patterns under the haploid (dashes) and diploid Li and Stephens model over a simulated reference set of *n* (horizontal axis) haploid sequences, using the Viterbi (red) and fastLS (*GR*een) algorithms, using *ρ*/*μ* = 2. Dots represent measurements, curves show quadratic fits. **a.** Results for a simulated reference population of *n* samples. **b.** Results for a fixed simulated reference population of 100,000, subsampled to *n* samples.

## 6 Performance

For testing the fastLS algorithms were implemented in *C*++, with all tables stored in uncompressed form in memory. To validate the implementations and to compare runtimes, standard Viterbi algorithms for the haploid and diploid LS model were also implemented. The fastLS algorithms included the traceback step, but Viterbi traceback this was le*β* out because of memory constraints. Two sets of simulations were performed. For the first, 30 Mb of sequence in populations of size 100 to 10, 000 were simulated by `scrm` [12] using the ’standard simulation’ model of [13]. For each population *i* simulated an additional 50 samples to serve as input sequences. This resulted in a number of segregating sites ranging from from 129, 945 for the 150-sample case, to 436, 361 for 10,050 samples. For the second set, I simulated a single population of 100,000 samples under the same model (resulting in 621, 156 segregating sites) and sub-sampled reference populations of 100 to 10, 000 samples from these.

The run-times of the Viterbi algorithms show the expected linear and quadratic dependence on *H*. The fastLS algorithms show a sub-linear dependence on *H*. In the case of the sub-sampled population, which have a fixed number of loci (not all of which segregate in the sample), the dependence on *H* is weakest, and in fact the diploid algorithm becomes faster for larger populations, probably because longer haplotype matches can be found in larger populations, resulting in more efficient pruning of the prefix sets.

## 7 Acknowledgements

Thanks to Sorina Maciuca and Zam Iqbal who introduced me to the idea of position symbols which led directly to this work; and to Gil McVean and Jerome Kelleher for helpful comments on the manuscript.

## 8 Appendix

### 8.1 Derivation of Algorithm 3

To derive the PBWT algorithm for sequence matching I first need some expressions that describe the structure of *M*. From Figure 1 it can be seen that

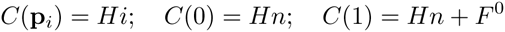

where 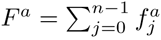 is the number of symbols *a* ∈ {0,1} in *x*, and 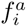 is the (haplotype) frequency of *a* at position *i* in *x*_0_,…,*x*_*H*‒1_. Let 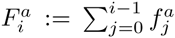 be the cumulative haplotype frequency across positions up to *i* – 1, and set 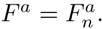. Then *R*^*a*^(*i*) satisfies

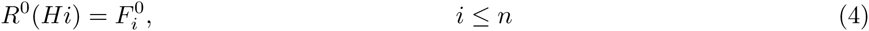

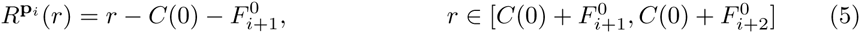

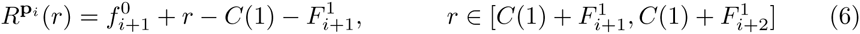

I define 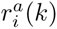 so that

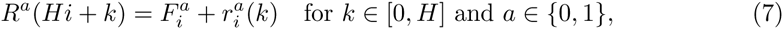

or equivalently, 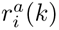 counts how often a appears in *BWT*(*X*)[*Hi*, *Hi* + *k*). To derive the PBWT sequence matching algorithm, it suffices to track one of the bounding variables, say *S*, for two steps through Algorithm 2. Assume that the subsequence matched so far starts at position *i*, so that *S* = *Hi* + *K* with *K* ∈ [0, *H*], and that the next character to be matched is *a* ∈ {0,1}. The first step replaces *S* with

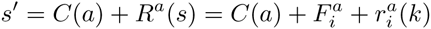

where the second equality follows from (7). The function 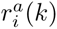 returns the number of occurrences of a before the *k*th row within the block starting at row *iH* in *M*. This block includes all sequences that start with **p**_*i*_, so that 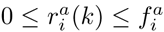 for *k* ∈ [0, *H*], and the conditions for (5) and (6) apply, allowing the result of the second step to be computed. The sequence now ends with the symbol **p**_*i*‒1_, so that if *a* = 0, *S*′ is replaced by

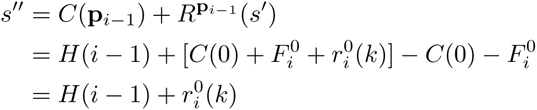

whereas if *a* = 1,

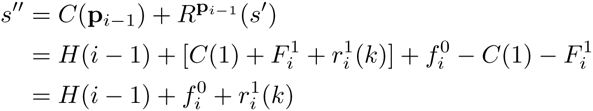

Since *R*^0^(*r*) + *R*^1^(*r*) = *r* for *r* ≤ *Hn*, it follows that 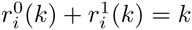 for 0 ≤ *k* ≤ *n*, so that the last-to-first function mapping *k* to the new value *k*′ satisfying *S*′′ = *H*(*i* – 1) + *k*′ is *LF*(*k*, *a*, *i*) as defined in (3).

### 8.2 Proof of Theorem 1

The key observation is that if *S*′ contains a path *π*′ with state (*G*′, *S*′), then *S*′ does not need to contain any path *π* (of the same length) with state (*G*, *s*) if *G*′ ⊇ *G* and *S*′ ≤ *S*. In this case I say that *π*′ *undercuts π*, or symbolically *π*′ ≤ *π*. In addition, if *π*′*R* ≤ *π* I also say that *π*′ ≤ *π*, again because all scores that are achievable with *π* as prefix are also achievable with prefix *π*′.

Since *S* is a full prefix set for *x*[0,*l*), a trivial full prefix set for *x*[0,*l* + 1) is formed by the union of simple extensions 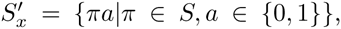, and recombination extensions 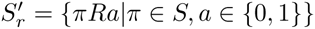. To prove that 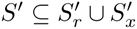 is also a full prefix set, we need to show that any path 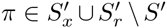 is undercut by some path *π*′ ∈ *S*′. In the proof below I will identify for any such *π* a *π*′ that *strictly* undercuts *π* (written as *π*′ < *π*) – that is, either the score is strictly lower or the group is strictly larger – but which is not necessarily an element of *S*′. If an element is found that is not in *S*′, the process can be repeated, finding a *π*′′ < *π*′ < *π*, and so forth. This process has to stop eventually, with an element in *S*^′^, because *S* cannot decrease indefinitely and *G* cannot increase indefinitely.

#### proof

First consider an arbitrary element 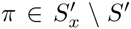. Because *π* ∉ *S*′ we have 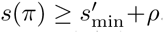. Consider *π*′*R* with *π*′ ∈ *S*′ such that 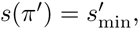, then 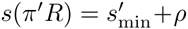 and *G*(*π*′ *R*) = Ω ⊃ *G*(*π*), so that *π*′*R* < *π*, and therefore *π*′ < *π*.

Next, consider an arbitrary element of 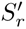, say *πR*a. We may assume that *s*(*π*) = *s*_min_, as otherwise *π*′*R*a with *s*(*π*′) = *s*_min_ strictly undercuts it. We may also assume that *a* = *x*[1], since otherwise let *πc* be some extension of *π* (which must exist), then *s*(*πcR*) ≤ *s*(*π*) + *μ* + *ρ* = *s*(*πRa*) and *G*(*πcR*) = Ω ⊃ *G*(*πRα*) so that *πcR* < *πRa* and therefore *πc* < *πRa*. Finally, if *πa* exists, then *s*(*πRa*) = *s*(*πRa*) and *G*(*πRa*) ⊃ *G*(*πRa*) so that *πa* < *πRa*. This completes the proof.

### 8.3 Proof of Theorem 2

The structure of this proof is identical to the previous one. The equivalent key observation is that a full prefix set *S*′ does not need to contain a path pair {*α,β*} if *S*′ already contains a path pair {*α*′,*β*′}with *s*(*α*′,*β*′) ≤ *s*(*α*,*β*)and *G*(*α*′,*β*′) ⊇ *G*(*α*,*β*);in this case I say that the path pair {*α*′, *β*′} undercuts {*α*, *β*}, or symbolically {*α*′,*β*′} ≤ {*α*,*β*}. I also write {*α*′,*β*′} ≤ {*α*,*β*} if any one of {*α*′*R*,*β*′} ≤{*α*,*β*}, {*α*′,*β*′*R*} ≤{*α*,*β*} or {*α*′*R*,*β*′*R*} ≤{*α*,*β*} is true.

A trivial full prefix set for *x*[0,*l* + 1) is formed by the union 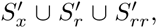, where *s*_*x*_ = {{*αa*, *βb*}|{*α*, *β*} ∈ *S*; *a*, *b* ∈ {0,1}}, and *S*_*rr*_ = {{*αRa*, *βRb*}|{*α*, *β*} ∈ *S*; *a*, *b* ∈ {0,1}}. The task is to prove that any path pair in 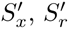 or 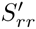 but not in *S*′ is undercut by some element of *S*′, and again I do this by identifying for any 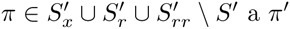 that strictly undercuts *π*.

#### proof

Consider an arbitrary 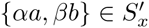 not in *S*′, so that 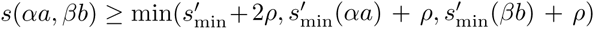. Suppose first that 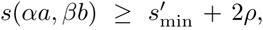, and let *α*′a′ and *β*′*b*′ be such that 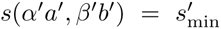, then *G*(*α*′*a*′*R*, *β*′*b*′*R*) ⊃ *G*(*αa*, *βb*) and 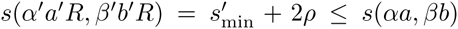, so {*α*′*a*′*R*,*β*′*b*′*R*} < {*αa*, *βb*}, and so {*α*′*a*′,*β*′*b*′} < {*αa*, *βb*}. Alternatively, suppose that 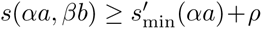, and let *β*′*b*′ be a path so that {*α*, *β*′} ∈ *S* and 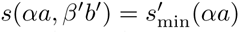, then *G*(*αa*, *β*′*b*′*R*) ⊃ *G*(*αa*, *βb*) and 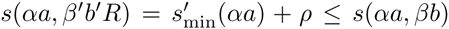, so that {*αa*, *β*′*b*′*R*} < {*αa*, *βb*}, and so {*αa*, *β*′b′} < {*αa*, *βb*}. The case 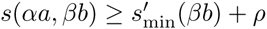 is dealt with similarly.

Next, consider an arbitrary element 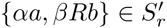. We may assume that *s*(*α*, *β*) = *s*_min_ (*α*) as otherwise it is possible to undercut this pair by choosing *β* appropriately. We may also assume that no {*α*, *β*′} exists in *S* so that *s*(*α*, *β*′) = *s*_min_(*α*) and **αa** and *β*′b are extensions, for if such a pair exists, the pair {*αa*, *β*′*bR*} undercuts {*αa*, *βRb*} as it achieves the same score and has a strictly larger group. Now suppose *x*[*l*] = 1. If *a* + *b* ≠ 1, for any extension *βb*′ of *β* we have *s*(*αa*, *βb*′*R*) ≤ *s*(*α*, *β*) + *μ* + *ρ* = *s*(*αa*, *βRb*) and *G*(*αa*, *βb*′*R*) ⊃ *G*(*αa*, *βRb*) so that {*αa*, *βb*′*R*} < {*αa*, *βRb*}, as required. To deal with the case *x*[*l*] ≠ 1, say *x*[*l*] = 0, suppose *b* = 1 and let *βb*′ be any extension, then *s*(*αa*, *βb*′*R*) ≤ *s*(*α*, *β*) + (*α* + 1)*μ* + *ρ* = *s*(*αa*, *βR*1) so that {*αa*, *βb*′*R*} < {*αa*, *βR*1}, as required. The case *x*[1] =2 is dealt with similarly.

Finally, consider an arbitrary element 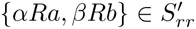. As before we may assume that *s*(*α*, *β*) = *s*_min_. Let’s first deal with the case *x*[1] = 1. If *a* = *b* then let *αa*′ be an arbitrary extension, then *s*(*αa*′*R*′, *βRb*) ≤ *s*(*α*, *β*) + *μ* + 2*ρ* = *s*(*αRa*, *βRb*) and *G*(*α*a*′R*, *βRb*) ⊃ *G*(*αRa*, *βRb*) so {*α*a′*R*, *βRb*} < {*αRa*, *βRb*}. If instead *a* ≠ *b*, then let *αa*′ and *βb*′ be arbitrary extensions. If *a*′ = *a* then {*αa*′*R*, *βRb*} < {*αRa*, *βRb*} by a now familiar argument. If *b*′ = *b* then {*αRa*, *βb*′*R*} is the required strictly undercutting path pair. If both *a*′ ≠ *a* and *b*′ ≠ *b* then *a*′ ≠ *b*′ and {*αa*′*R*, *βb*′*R*} achieves the same score and a larger group, and therefore strictly undercuts {*αRa*, *βRb*}. It remains to deal with the case *x*[*l*] ≠ 1, say *x*[*l*] = 0. If either *a* = 1 or *b* = 1 (or both), say *b* = 1, then let *βb*′ be an arbitrary extension, then *s*(*αRa*,*βb′R*) ≤ (*a* + 1)*μ* + 2*ρ* = *s*(*αRb*, *βRb*) so that {*αRa*,*βb′R*} < *αRa*,*βRb*}. So we can assume that *a* = *b* = 0. The argument in the case *x*[*l*] = 2 is similar. Finally, suppose there is a pair {*α*′, *β*′} with *s*(*α*′, *β*′) = *s*_min_ and either *α*′*b* or *β*′*b* is an extension, say *β*′*b* is, then {*αRb*,*Rb*, *β*′*Rb*} ≤ {*αRb*, *βRb*} as required. This completes the proof.

